# Sub physiological temperature induces pervasive alternative splicing in Chinese hamster ovary cells

**DOI:** 10.1101/863175

**Authors:** Ioanna Tzani, Craig Monger, Krishna Motheramgari, Clair Gallagher, Ryan Hagan, Paul Kelly, Alan Costello, Justine Meiller, Patrick Floris, Lin Zhang, Martin Clynes, Jonathan Bones, Niall Barron, Colin Clarke

**Author notes:** Equal contribution (Ioanna Tzani, Craig Monger). **Abbreviations:** Alternative splicing (AS); Chinese hamster ovary (CHO); monoclonal antibody (mAb); next generation sequencing (NGS); RNA sequencing (RNASeq); non-sense mediated decay (NMD); temperature shifted (TS); non-temperature shifted (NTS); Benjamini Hochberg (BH); skipped exon (SE); mutually exclusive exon (MXE); untranslated region (UTR); retained intron (RI); alternative 5’ and 3’ splice sites (A5SS and A3SS); percent spliced in (Δψ).

## Abstract

RNA sequencing (RNASeq) has been widely used to associate alterations in Chinese hamster ovary (CHO) cell gene expression with bioprocess phenotypes, however alternative mRNA splicing, has thus far, received little attention. In this study, we utilised RNASeq for transcriptomic analysis of a monoclonal antibody producing CHOK1 cell line subjected to a temperature shift. More than 2,365 instances of differential splicing were observed 24hrs after the reduction of cell culture temperature. 1,163 of these alternative splicing events were identified in genes where no changes in abundance were detected by standard differential expression analysis. Ten examples of alternative splicing were selected for independent validation using qPCR in the monoclonal antibody producing CHOK1 line used for RNASeq and a further 2 CHOK1 cell lines. This analysis provided evidence that exon skipping and mutually exclusive splicing events occur in genes linked to the cellular response to changes in temperature and mitochondrial function. While further work is required to determine the impact of changes in mRNA sequence on cellular phenotype, this study demonstrates that alternative splicing analysis can be utilised to gain a deeper understanding of post-transcriptional regulation in CHO cells during biopharmaceutical production.

## Introduction

Chinese hamster ovary (CHO) cells are the principal expression host for the manufacture of recombinant therapeutic proteins with 84% of all monoclonal antibodies (mAb) products now produced in this cell line (Walsh, 2018). For more than a decade, academic and industrial research groups have sought to improve our knowledge of these cell factories and the molecular characteristics that underpin the efficient production of biopharmaceuticals (Kuo et al., 2018). The goal of these efforts is to complement advances in traditional upstream manufacturing technologies and further intensify production efficiencies by leveraging understanding of the CHO cell biological system to select and/or rationally design ultra-high-performance cell lines (Fischer, Handrick, & Otte, 2015). The pace of research in this area has accelerated following the development of next generation sequencing (NGS) technology. The first genome sequence for the CHOK1 genome was published in 2011 (Xu et al., 2011) and there are now multiple nuclear as well as mitochondrial genome sequences available publicly for CHO cell lines and the Chinese hamster (Brinkrolf et al., 2013; Kaas, Kristensen, Betenbaugh, & Andersen, 2015; Kelly et al., 2017; Lewis et al., 2013; Rupp et al., 2018).

In particular, our ability to analyse the CHO cell transcriptome has improved markedly in the post-genomic era (Monger et al., 2015). RNA sequencing (RNASeq) approaches have enabled the accurate quantitation of variations in gene expression associated with critical industrial phenotypes such as cellular growth rate, cell specific productivity (Sha, Bhatia, & Yoon, 2018) and product quality (Clarke et al., 2019). The application of RNASeq has also played a pivotal role in the identification of promoter regions in the CHO cell genome, improving the annotation of small non-coding RNAs (Gerstl, Hackl, Graf, Borth, & Grillari, 2013; Hackl et al., 2012) and opening new routes for genetic engineering to improve CHO cell line performance (Raab et al., 2019). Although tremendous advances in characterization of the CHO cell RNA accomplished using NGS technology have been made in recent years, previous studies have largely focused on correlating the abundance of RNA with bioprocess traits in CHO cells, much in the same way the field utilised traditional microarray technology - as others have noted this bias toward using RNASeq to identify variation in expression levels is not confined to CHO cell biology (Hartley & Mullikin, 2016). To advance our understanding of CHO cell biology we must maximize the utility of RNASeq to provide not only accurate quantitation, but also focus on associating differences in mRNA sequence composition with CHO cell behaviour and performance in biopharmaceutical manufacturing.

The removal of introns and subsequent ligation of exons from a pre-mRNA molecule to produce a mature mRNA is known as splicing. In eukaryotic organisms, a protein-RNA machine known as the spliceosome (Will & Lührmann, 2011) coordinates the selection of different exon and splice sites to enable the creation of distinct mRNA isoforms from a single gene. This process, termed alternative splicing (AS), is recognized as a central step in the regulation of gene expression in eukaryotic cells responsible for the remarkable transcriptome and proteome diversity arising from a limited set of genes (Pan, Shai, Lee, Frey, & Blencowe, 2008; Wang et al., 2008). The production of different mRNA isoforms from a single gene via alternative splicing can result in a variety of functional impacts ranging from subtly altering the activity of the protein to global regulation of biological processes (Kelemen et al., 2013). For instance, a isoform of *Xpb1* is required for the activation of the unfolded protein response (Yoshida, Matsui, Yamamoto, Okada, & Mori, 2001). Alternative splicing is well known to play an important role in apoptosis with different splice variants of the *BCL2L1* gene encoding proteins reported to have both pro- and anti-apoptotic roles (Boise et al., 1993). Alternate splicing can also direct proteins to different cellular compartments or control membrane binding (e.g. removal of an exon of the Cyclin D1 gene results in retention of the mRNA in the nucleus (Lévêque, Marsaud, Renoir, & Sola, 2007)). In addition, AS can also regulate gene expression through the production of non-functional proteins via the incorporation of premature stop codons resulting in non-sense mediated decay (NMD) (Ge & Porse, 2014).

A strategy commonly referred to as a “temperature shift” is often utilized to improve the performance and extend the lifetime of mammalian cell culture processes in the biopharmaceutical industry (Masterton & Smales, 2014). In this study, we examined the transcriptomic response to a temperature shift using high resolution RNASeq to not only detect differences in expression at the gene level, but also, for the first time in CHO cells, monitor significant variations in mRNA splicing. Through this analysis we have uncovered hundreds of significant alterations in exon and splice site usage where no change in expression could be identified using gene-level differential expression analysis. In addition, we examined the occurrence of a cohort of differential splicing events identified by RNASeq in 3 CHO cell lines using qPCR. The findings of this study reveal a previously unstudied aspect of the CHO cell’s molecular response to sub-physiological cell culture temperature and illustrate the limitations of focusing solely on gross changes in gene expression during transcriptomic analysis of CHO cells.

## Methods

### Cell culture and sample preparation

A mAb producing CHOK1 cell line (CHOK1-mAb) was seeded at a density of 2 × 10^5^ cells/ml in SFM-II media (Gibco, 12052098) in a Kuhner orbital shaker (170rpm) at 5% CO_2_. Eight replicate shake flask cultures were grown at 37°C for 48hr post-seeding; at this point the temperature of 4 shake flasks was reduced to 31°C (referred to as the temperature shift (TS) sample group), while the remaining 4 shake flasks were maintained at 37°C (referred to as the non-temperature shift (NTS) sample group). 5 × 10^6^ cells were harvested from each shake flask at 72hrs post-seeding and RNA extracted for transcriptomics using TRIzol (Thermo Fischer Scientifc, MA) (Figure 1A).

**Figure 1:**
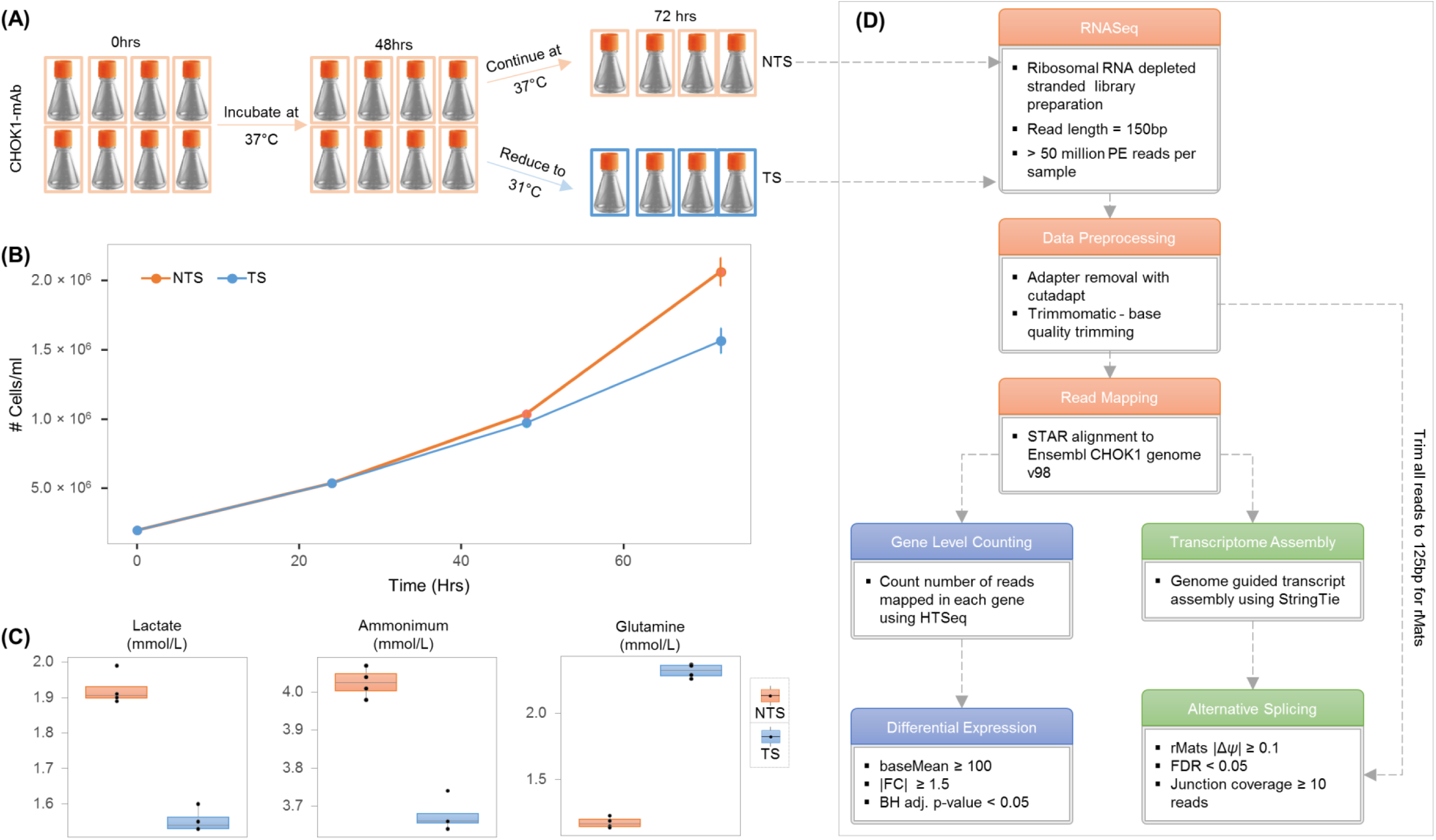
Reducing cell culture temperature decreases the growth rate of the CHOK1-mAb cell line and alters the concentration of extracellular metabolites. **(A)** 8 biological replicates were seeded at 2 × 10^5^ cells/ml. At 48hrs post seeding the temperature of 4 of the replicates was reduced from 37°C to 31°C. Total RNA was extracted from the cultures at 72hrs post seeding and 24hrs post temperature shift. **(B)** A significant reduction in cell counts between the NTS and TS sample groups was observed 24hrs post temperature shift. **(C)** Metabolite profiling of the cell culture media identified a significant increase in glutamine concentration as well as a reduction in lactate and ammonia in temperature shifted samples. **(D)** Following ribosomal RNA depletion a strand specific library was prepared for RNASeq. Data was preprocessed and aligned to the CHOK1 genome. At this point 2 separate strategies were used to analyses the data: (1) a standard gene-level count based differential expression analysis using the v95 Ensembl annotations. (2) A genome guided assembly was constructed using the CHOK1 genome using the RNASeq data and the Ensembl CHO1K genome followed by the utilisation of rMats to identify differential splicing.

### Metabolite and pH analysis

A Nova Flex (Nova Biomedical, Waltham, MA) was used to measure the concentration of glutamine, glutamate, glucose, lactate, ammonium, calcium, sodium and potassium in the media for the NTS and TS sample groups at 72hrs post-seeding. Changes in metabolite concentrations between the groups was considered statistically significant if the difference in means yielded a p value < 0.05 using a two-tailed t-test.

### RNASeq library preparation

Total RNA was processed for library construction as follows: Illumina RiboZero Gold rRNA-probes (Illumina, San Diego, CA) were hybridized to total RNA to remove nuclear-encoded and mitochondrial ribosomal RNA (rRNA). The rRNA depleted samples were fragmented and sequencing libraries prepared using the TruSeq Stranded Total RNA Library Prep Kit (Illumina, San Diego, CA). Briefly, first-strand cDNA synthesis was performed using reverse transcriptase and random primers in the presence of Actinomycin D, followed by second-strand cDNA synthesis with DNA polymerase I and RNase H. Double-stranded cDNA was end-repaired and A-tailed for subsequent adapter ligation. Indexed adaptors were ligated to the A-tailed cDNA. Enrichment by PCR was performed to generate the final cDNA sequencing library. Libraries were sequenced on an Illumina NextSeq500 (Illumina, San Diego, CA) configured to yield at least 50 million 150bp paired-end reads per sample.

### Data pre-processing and read mapping

Adapters were removed from the raw RNA sequencing reads using cutadapt v1.18 (Martin, 2011) before quality trimming was performed using Trimmomatic v0.36 (Bolger, Lohse, & Usadel, 2014). Preprocessed reads were subsequently aligned to the Ensembl v98 CHOK1 reference genome sequence using the STAR v2.7.2d (Dobin et al., 2013) in combination with splice junction information from the Ensembl CHOK1 reference annotation to increase read mapping rates. The mapping accuracy was assessed using RNA-SeQC v1.1.8 (DeLuca et al., 2012) for each of the 8 samples.

### Differential Gene Expression Analysis

To determine the number of genes differentially expressed between the NTS and TS sample groups the number of reads mapping to protein coding genes was counted with HTSeq (Anders, Pyl, & Huber, 2015). The DESeq2 Bioconductor package (Love, Huber, & Anders, 2014) was utilised to analyse differential gene expression between the NTS and TS sample groups. Gene expression levels were normalised for sequencing depth using DESeq2, with gene expression changes of at least ±1.5 fold with an accompanying Benjamini Hochberg (BH) adjusted p-value of <0.05 considered significantly differentially expressed.

### Genome guided assembly and analysis of alternative splicing

To increase the number of mRNA isoforms annotated for alternative splicing analysis a genome guided assembly was constructed using StringTie v2.0.3 (Pertea et al., 2015) (minimum junction coverage = 5). Spurious transcripts that arose when two genes had same-strand overlap were removed the from StringTie annotation file (the two transcripts were merged into a single transcript in this case). The *gffcompare* tool was utilised to compare the StringTie transcriptome assembly to the CHOK1 Ensembl reference annotation.

### Identification of alternative splicing events

rMats v4.0.2 (Shen et al., 2014) was used to identify differential alternative splicing using the read alignments and StringTie annotation file. Prior to analysis, preprocessed sequencing reads were trimmed to 125bp using the *trimFastq*.*py* tool provided as part of the rMats suite. The rMats statistical model requires that all input reads are the same length. Differential alternative splicing events were considered to be significant if the absolute change in “percent spliced in” (denoted as Δ*ψ*) was ≥ 0.1 with a FDR < 0.05 and at least 10 aligned reads spanning the exon junctions.

### Enrichment analysis

The GOrilla tool (Eden, Navon, Steinfeld, Lipson, & Yakhini, 2009) was utilised to determine if GO terms were over-represented in the genes found to be differentially expressed and those genes found to be exclusively regulated via alternative splicing. An enrichment FDR q-value < 0.05 was considered statistically significant.

### Reproducibility of the RNASeq data analysis

The raw RNA sequencing data has been deposited to NCBI SRA (PRJNA593052) and the code required to reproduce the differential expression and differential splicing analysis will be released at https://github.com/clarke-lab/cho_cell_AS upon acceptance of this manuscript.

### qPCR validation of selected alternative splicing events

To validate the alternative splicing events using qPCR, the CHOK1-mAb cell line was again (independent to the samples for RNASeq) cultured as described above. To determine if alternative splicing events could be identified in additional CHO cell cells, a non-producing CHOK1 cell line acquired from ATCC (CHOK1-ATCC) along with a second CHOK1 cell line producing an anti-IL8 mAb (CHOK1-DP12) were also analysed by qPCR. The CHOK1-ATCC and CHOK1-DP12 cell lines were cultured using the same model of temperature shift previously described however these cells were maintained in BalanCD CHO Growth A Medium (Irvine Scientific, 91128) supplemented with 6mM and 8mM L-glutamine respectively.

Total RNA was extracted from cell pellets using Trizol (Invitrogen, cat. no 15596026) (Sigma AMPD1) according to the manufacturers’ specifications. The quality of RNA was assessed by electrophoresis on 1% agarose gels in TAE where all samples showed bands typical of undegraded RNA and the purity as well as quantity of RNA was assessed by spectrophotometry (Nanodrop 1000; Thermo Scientific). cDNA was generated from 2μg of total RNA using the Applied Biosystems kit (ref 4368814), according to the manufacturer’s guidelines.

Ten alternative splicing events identified by rMats were selected for qPCR validation in the three cell lines representing skipped exons (SE) (*Immp1l, Mff, Slirp, Gpx8, Dnm1l, Dnajc15 and Hikeshi*) and mutually exclusive exons (*Rnf34, Fars2* and *Ubr2*) (MXE) events found to have significant positive or negative Δ*ψ* between the NTS and TS groups. We also validated differences in the abundance of the *Cirbp* gene using qPCR to determine the difference in gene expression between the two groups. Primers were designed to span splice junctions whenever possible, using the default setting of PrimerQuest Real-Time PCR Primer Design Tool (IDT). Primer-Blast (Ye et al., 2012) was used to assess *in silico* primer specificity, self-complementarity and 3’ self-complementarity. Negative controls containing no reverse transcriptase and water controls were included in each qPCR experiment to monitor potential contamination with gDNA or in the reaction set up respectively. Primer specificities were assessed by agarose gel electrophoresis of qPCR products and melting curve analysis at the end of the qPCR runs. Each qPCR reaction was carried out in a 20μl final volume containing 4μl of 1:10 diluted cDNA, 10 μl of the Fast SYBR Green Master Mix (Applied BioSystems, cat.no 4385616), 0.8 μl of 10uM for each forward and reverse primer (400 nM final concentration) and 4.4 μl of nuclease free water. Following an initial denaturation step at 95°C for 20s, temperature cycling was performed for a total of 40 cycles. Each cycle consisted of denaturation at 95°C for 3s and annealing and extension at 60°C for 30s. Each qPCR reaction was performed in triplicate per sample (n=3) in an AB7500 Real Time PCR instrument. Ten-fold serial dilutions of gBlock fragments (Integrated DNA Technologies, IDT) were used to generate standard curves between log DNA copies (10^2^ to 10^8^ copies) and quantification cycle (C_q_) values. The standard curves were used both for absolute quantification of mRNA and measurement of amplification efficiency with the equation E =10(−1/slope). The number of RNA copies in each sample was calculated from the following equation: RNA copies = 10(C_q_-b)/slope, where b is the y-intercept. A ≥ 10% difference along with a two-tailed t-test p-value < 0.05 difference in the number of RNA copies between the NTS and TS groups was considered significant.

## Results

### Reduction of cell culture temperature decreases growth rate and alters the extracellular metabolite profile of the monoclonal antibody CHOK1 cell line

To determine the effect of a temperature shift on a mAb producing CHO cell line (CHOK1-mAb), 8 replicate shake flasks were grown for a period of 72hrs. For those samples designated as the control group (n=4), referred to as non-temperature shifted (NTS), the incubator temperature was maintained at 37°C for the duration of cell culture. The second sample group (n=4), referred to as temperature shifted (TS), was subjected to a reduction in temperature from 37°C to 31°C at 48hrs post-seeding (Figure 1A) and cultured for a further 24hrs. The cell counts for the 8 cultures replicates remained comparable prior to temperature reduction at 48hrs, however at the 72hr timepoint the cells/ml in the TS sample group was 24% lower that of the NTS samples (Figure 1B, Table S1A).

Profiling of the cell culture media using a Nova Flex analyser was carried out to assess the impact of reducing the cell culture temperature on extracellular metabolites (Table S1B, Figure S1). The glutamate concentration of the TS sample group decreased by 3% and although the average glucose concentration was found to be 20% higher in the TS sample group, this result was not statistically significant. The concentrations of ammonia and lactate were reduced by 8% and 19% respectively in the TS sample group while the concentration of glutamine increased by 97% (Figure 1C, Table S1B).

### Ribosomal RNA depleted, strand specific library preparation yields high quality libraries for CHO cell RNASeq

Following extraction of total RNA from the NTS and TS sample groups at 72hrs post-seeding, the samples were rRNA depleted prior to strand-specific library preparation for Illumina sequencing (Figure 1D). Between 50 million and 77 million paired end reads were sequenced for each of the 8 samples and upon completion of data pre-processing the number of paired end reads remaining ranged from 42 million to 66 million (Table S2). An average of 86% of the surviving read pairs were confidently mapped against the Ensembl CHOK1 reference sequence using the STAR algorithm (the lowest proportion of mapped reads for a single sample was 84%). Of the total number of reads sequenced at least 72% could be aligned to the CHOK1 the genome for a given sample (Figure S2).

### Decreasing cell culture temperature induces widespread CHO cell gene expression changes

To identify “gene-level” changes in between the NTS and TS sample groups a count based differential expression analysis was conducted (Figure 1D). The HTSeq-count tool was used to determine the number of aligned reads within regions of the genome defined as protein coding using the Ensembl CHOK1 reference annotation. The resulting counts for each gene were normalised for differences in library size between samples before differential expression was carried out using DESeq2. Those genes which were altered in the TS samples by an absolute fold change ≥ 1.5 along with a BH adjusted p-value of < 0.05 were considered to be significantly differentially expressed. In total, 502 protein coding genes were found to upregulated, while 815 were found to be decreased in those samples subjected to the temperature shift (Figure 2A, Table S3)

**Figure 2:**
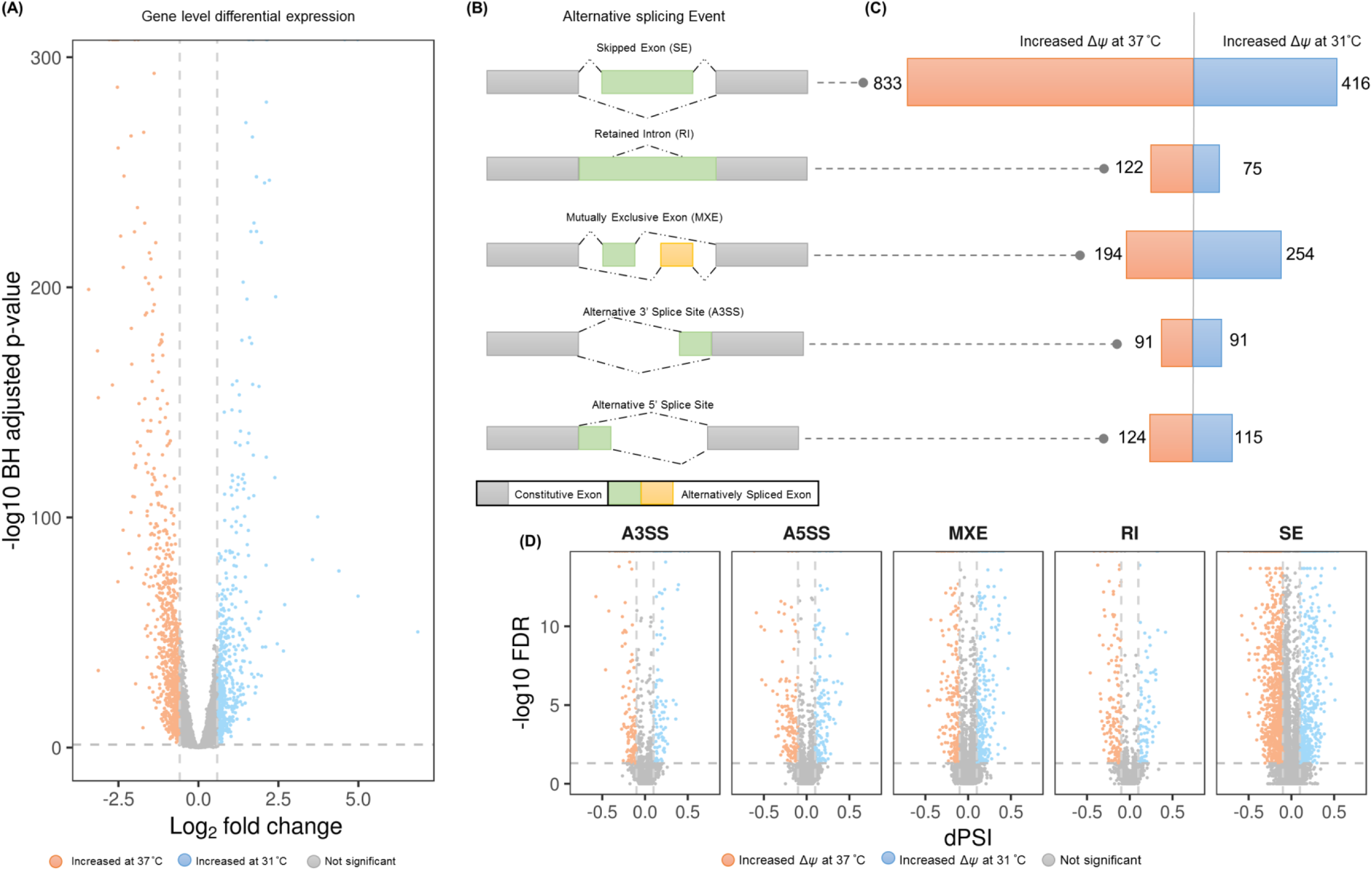
Reducing cell culture temperature induces differential gene expression and differential splicing in the CHOK1-mAb cell line. RNASeq and DESeq2 analysis revealed that (A) changes in gene expression with 1317 genes significantly increased or decreased after temperature shift. The rMats algorithm was utilized in combination with the Stringtie transcriptome assembly to determine if (B) 5 types of alternative splicing event – skipped exon (SE), mutually exclusive exon (MXE), retained intron (RI) as well as alternative 5’ and 3’ splice sites (A5SS and A3SS respectively). In total 2,365 alternative splicing events were identified as significantly differentially spliced (C) between the two sample groups with (D) Δψ values ranging from −0.736 to 0.546.

Examination of the genes differentially expressed revealed the presence of genes encoding well-known transcriptional indicators of CHO cell response to temperature shift including the cold inducible RNA binding protein (*Cirbp*) and RNA binding protein 3 (*Rbm3*) genes both significantly upregulated by 2.3 and 2.99 fold respectively in the TS sample group (Table S3).

### Expanding the Ensembl CHOK1 transcriptome annotation through genome guided assembly

To refine the current annotation of the Ensembl CHOK1 genome annotation prior to alternate splicing analysis a genome guided transcriptome assembly was performed using Stringtie. An individual StringTie transcriptome assembly was first constructed for the RNASeq data acquired from the NTS and TS groups before merging the individual assemblies and the current Ensembl annotation into a single unified assembly. Spurious transcripts that arose due to sense overlap of genes were removed from the StringTie assembly. Following comparison of the StringTie transcript annotation assembly to the current Ensembl annotation using the *gffcompare* tool an additional of 13,631 additional multi-exonic transcripts and >23,000 novel exons were identified.

### Widespread differential splicing is associated with a reduction of cell culture temperature

Differential splicing between the NTS and TS sample groups was carried out using the combined Ensembl reference and StringTie transcript annotation. The rMats algorithm requires input reads of the same length and prior analysis all pre-processed sequencing reads were trimmed to 125bp before being mapped to the CHOK1 genome using STAR. Utilisation of the StringTie transcriptome annotation was found to have increased the proportion of reads aligned to exons by 14% and more than doubled the number of transcripts detected (>5 exon mapping reads) in comparison to using the Ensembl CHOK1 reference genome alone (Figure S3). Differential splicing events identified by rMats were considered significant if an increase or decrease in the percent spliced in (*Δψ*) value ≥ 0.1, a FDR < 0.05 and at least 10 reads spanning the splice junction. In total, 2,365 significant differential splicing events were identified following comparison of the NTS and TS sample groups comprising skipped exons (n=1,299), mutually exclusive exons (n=448), retained introns (n=197) as well as alternative 5’ (n=239) and 3’ (n=182) splice sites were found to be associated with a decrease in cell culture temperature (Figure 2B, 2C and 2D, Table S4).

Following comparison of the 1,317 significantly differentially expressed genes to the 2,365 alternative splicing events identified using rMats, 1,163 genes were found to be differentially spliced but not differentially expressed between the NTS and TS sample groups (Figure 3A).

**Figure 3:**
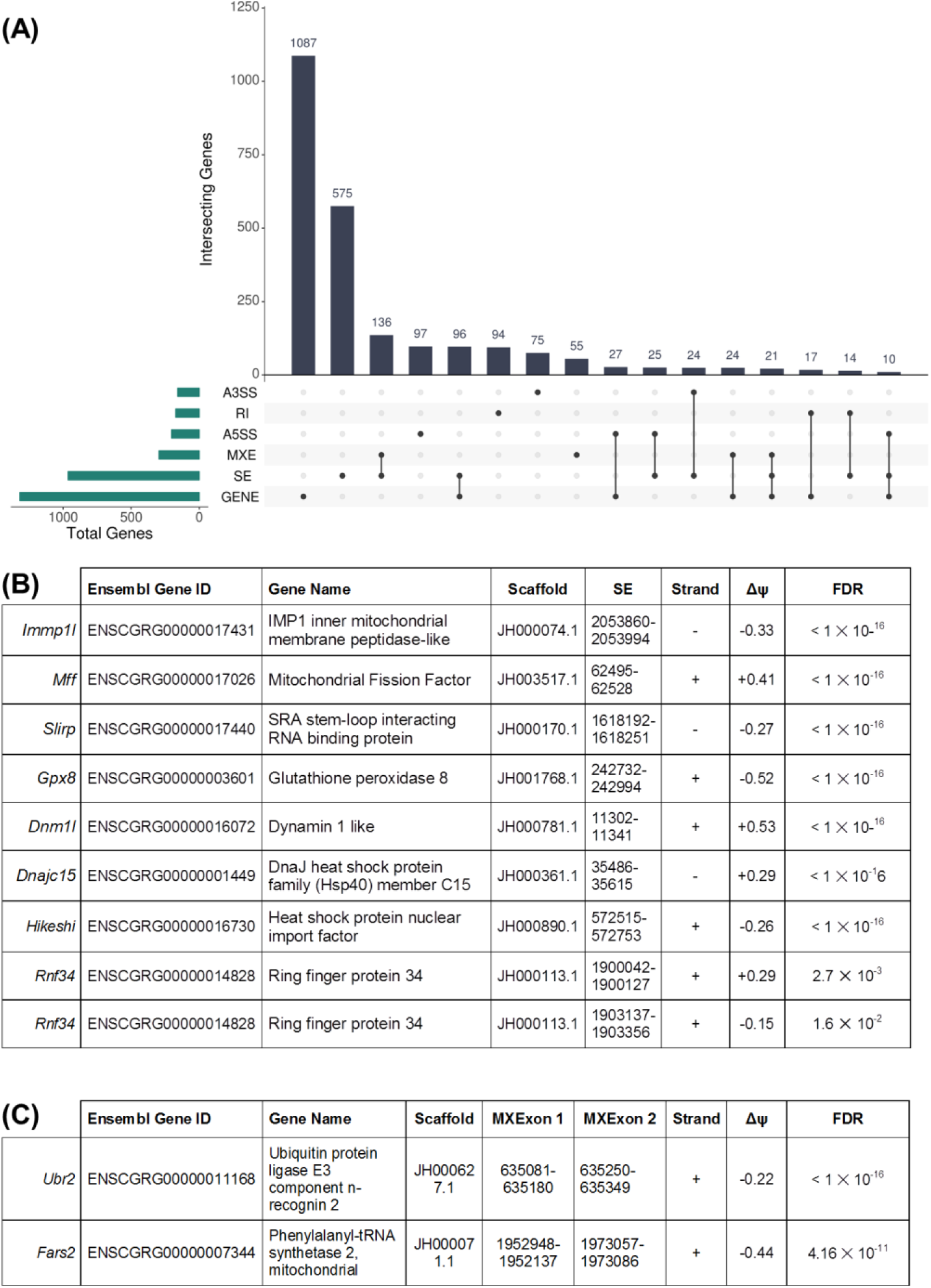
Comparison of gene-level differential expression and alternative splicing identifies genes regulated primarily through RNA splicing. **(A)** Those genes found to be differentially expressed at the gene-level or to be alternatively spliced between the NTS and TS conditions were compared and the results displayed using an UpSet plot. The lower portion of the UpSet plot displays all overlaps containing at least one gene from the 6 categories of transcriptional change i.e. differentially expressed genes (“GENE”), skipped exon (SE), mutually exclusive exon (MXE), retained intron (RI), alternative 5’ and 3’ splice sites (A5SS and A3SS respectively). A single black circle denotes the presence of the single transcriptional event corresponding to that category. Where one or more black circles are linked with a vertical black line there are overlapping genes between these categories. The bar graph directly above indicates the number of genes present. A cohort of 8 **(B)** skipped exon and **(C)** mutually exclusive splicing events identified using RNASeq that increased (blue) or decreased (orange) upon a reduction of cell culture temperature in the CHOK1-mAb cell line were chosen for further evaluation with absolute qPCR. *The 2 Rnf34 SE events found in neighbouring exons were anti-correlated and we reasoned that these changes in splicing could indicate the presence of a MXE event and designed PCR primers to target the possibility.

Enrichment analysis of the differentially expressed genelist was conducted using GOrilla against GO terms (biological processes, molecular function & cellular compartment) and significant overrepresentation of biological process categories including those related to cell cycle (GO:0007049) and cellular response to stress (GO:0033554) were identified (Table S5). Enrichment analysis against GO categories was carried out also for the 1,163 genes that are regulated exclusively through alternative splicing. Those categories found to be overrepresented included cell cycle (GO:0007049), response to cell stress (GO:0007049), as well as metabolic processes related to phosphate (GO:0007049). In addition, an enrichment of genes associated with the Golgi apparatus (GO:0005794) and mitochondria (GO:0005739) enriched within the splicing-only genelist.

### The CHOK1-ATCC and CHOK1-DP12 cell lines exhibit a variable response to temperature shift

An untransfected CHOK1 cell line acquired from ATCC (CHOK1-ATCC) and a CHOK1-DP12 cell line producing an anti-IL8 mAb were subjected to an identical cell culture model of temperature shift. A 10% reduction in cell number for CHOK1-ATCC cell line was observed following a reduction of cell culture temperature although this result was not statistically significant. The concentrations of glutamine and ammonia, however, were found to be 11% and decreased by 12% higher respectively (Table S1). For the CHOK1-DP12 cell line, cell numbers were 22% lower 24hrs post temperature shift, comparable to that of the CHOK1-mAb cell line, however no significant changes in extracellular metabolite concentrations could be identified (Table S1). Although there was variation in cell number and metabolite profiles, a comparable increase in *Cirbp* gene expression was found by qPCR indicating that temperature shift induced a transcriptional response in all 3 cell lines (Figure 4A, Table S8).

**Figure 4:**
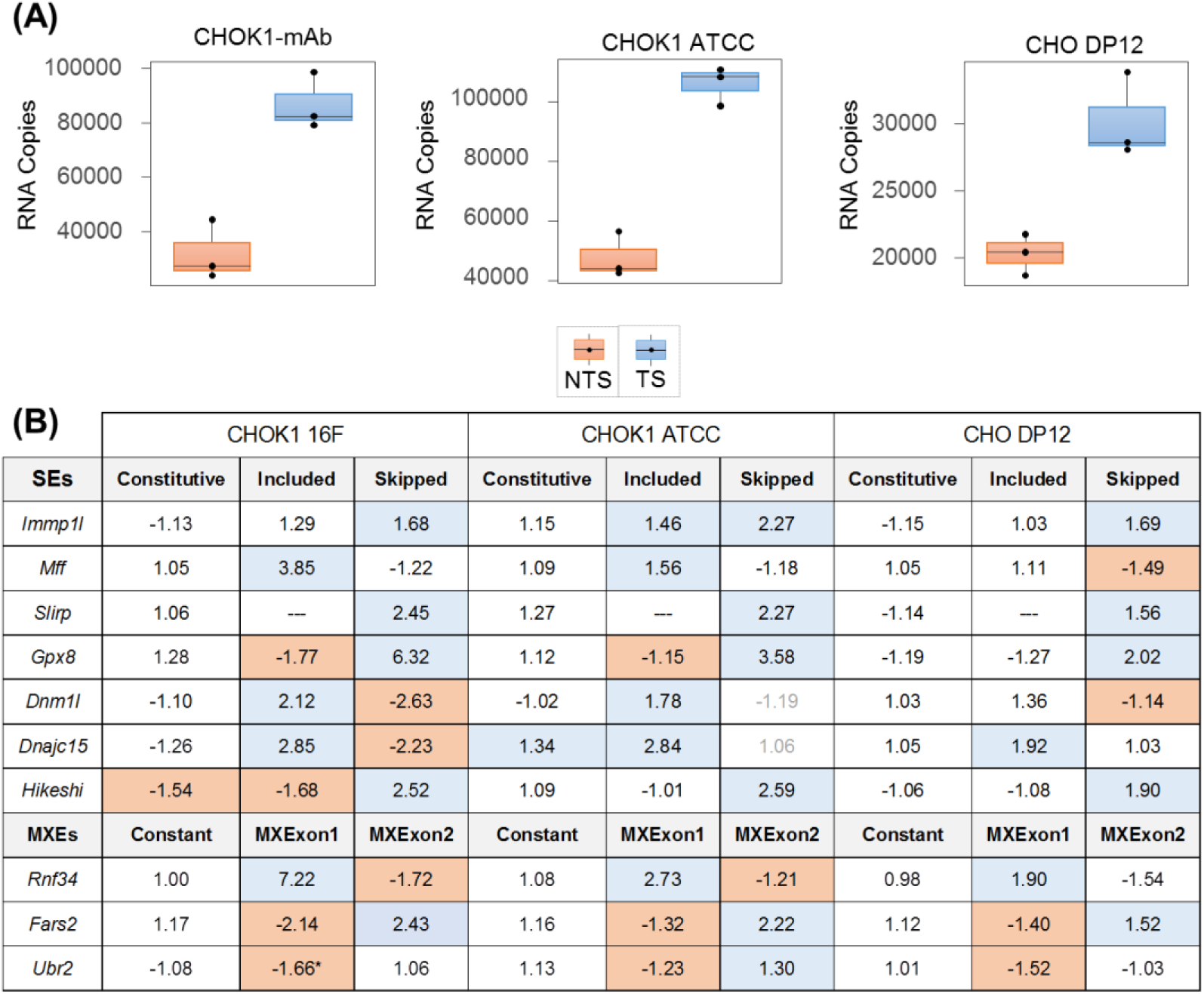
qPCR analysis of alternative splicing. The CHOK1-mAb cell line along with a CHOK1 cell line from ATCC and the CHOK1-DP12 cell line were subjected to the model of temperature shift utilized to generate the RNASeq data. For these samples a decrease in cell numbers as well as a significant increase in **(A)** *Cirbp* expression was observed at 24hrs post-temperature reduction. **(B)** Primers were designed to target those transcripts that contained either skipping or inclusion of an exon or one of the pairs of mutually exclusive exons. For the *Slirp* gene a specific primer could not be designed for the exon inclusion event. The largest degree of agreement between the RNASeq and qPCR was for the CHOK1-mAb while a lower degree of agreement was observed for the CHOK1-ATCC and CHOK1-DP12 cell lines. At least 1 of the 2 predicted events (skip/inclusion of exon1/exon2) was confirmed for all genes in at least one cell line. For each gene qPCR was also used to measure the expression of a constitutive exon (i.e. no alternative splicing detected using RNASeq). For this analysis a differential expression threshold of 1.5 fold in either direction with a p-value of < 0.05 was used similar to the thresholds used for DESeq2 algorithm. From this analysis only *Hikeshi* was found to be differentially expressed following the observation of a - 1.58 fold decrease (p = 1.3 × 10^−4^), in abundance in the temperature shifted samples.

### qPCR analysis of alternative splicing events associated with temperature shift in multiple CHO cell lines

A cohort of 10 alternative splicing events comprised of both SEs (Figure 3B) and MXEs (Figure 3B) identified in genes that were not differentially expressed was chosen for confirmation using an orthogonal method. These genes were selected due to their association with the cellular response to changes in temperature or role in mitochondrial function (due the observation of significant enrichment of the mitochondrion GO term for the splicing-only genelist). Following examination of the 2 *Rnf34* SE events (occurring in neighbouring exons) we found that the changes in *Δψ* were anti-correlated changes in exon skipping. Based on this evidence we reasoned that these changes in splicing could be a MXE and designed PCR primers to target the possibility.

The cell culture model of temperature shift was repeated for the CHOK1-mAb cell line and absolute real-time PCR used to assess changes in mRNA splicing (Table S6, Table S7, and Figure S4). The selected alternative splicing events were also examined in the NTS and TS sample groups from the CHOK1-ATCC and CHOK1-DP12 cell lines using qPCR. In addition, a constitutive exon (i.e. an exon not found to be alternatively spliced by rMats) was chosen for each gene (the only significant difference in expression observed was a downregulation of the *Hikeshi* constitutive exon in the CHOK1-mAb cell line (Figure 4B, Table S8).

For each of the selected SE events in 7 genes (*Immp1l, Mff, Slirp, Gpx8, Dnm1l, Dnajc15, Hikeshi*), two primer sets were designed to (1) amplify isoforms where the exon of interest was skipped (i.e. spliced-out) and (2) detect isoforms where the exon of interest was included (i.e. spliced-in). For each SE event the change in abundance of the “reciprocal event” was also determined (i.e. the expression of isoforms with the exon of interest spliced-in were also measured). Specific primers could not be designed to detect isoforms with the selected *Slirp* exon spliced in.

The temperature shift induced exon skipping increase in isoforms of the *Immp1l, Slirp, Gpx8* and *Hikeshi* genes discovered by RNASeq were confirmed by qPCR in all 3 CHO cell lines (Figure 4B, Table S8). A reciprocal decrease in inclusion of the *Gpx8* exon was observed in the two CHOK1 cell lines but not CHOK1-DP12 (Figure 5). *Hikeshi* isoforms containing the spliced in exon decreased in expression in CHOK1-mAb while no change in abundance was found in the other two cell lines. *Immp1l* exon inclusion increased in the CHOK1-ATCC cell line, however there was no significant change in expression for the CHOK1-mAb or CHOK1-DP12 cell lines.

**Figure 5:**
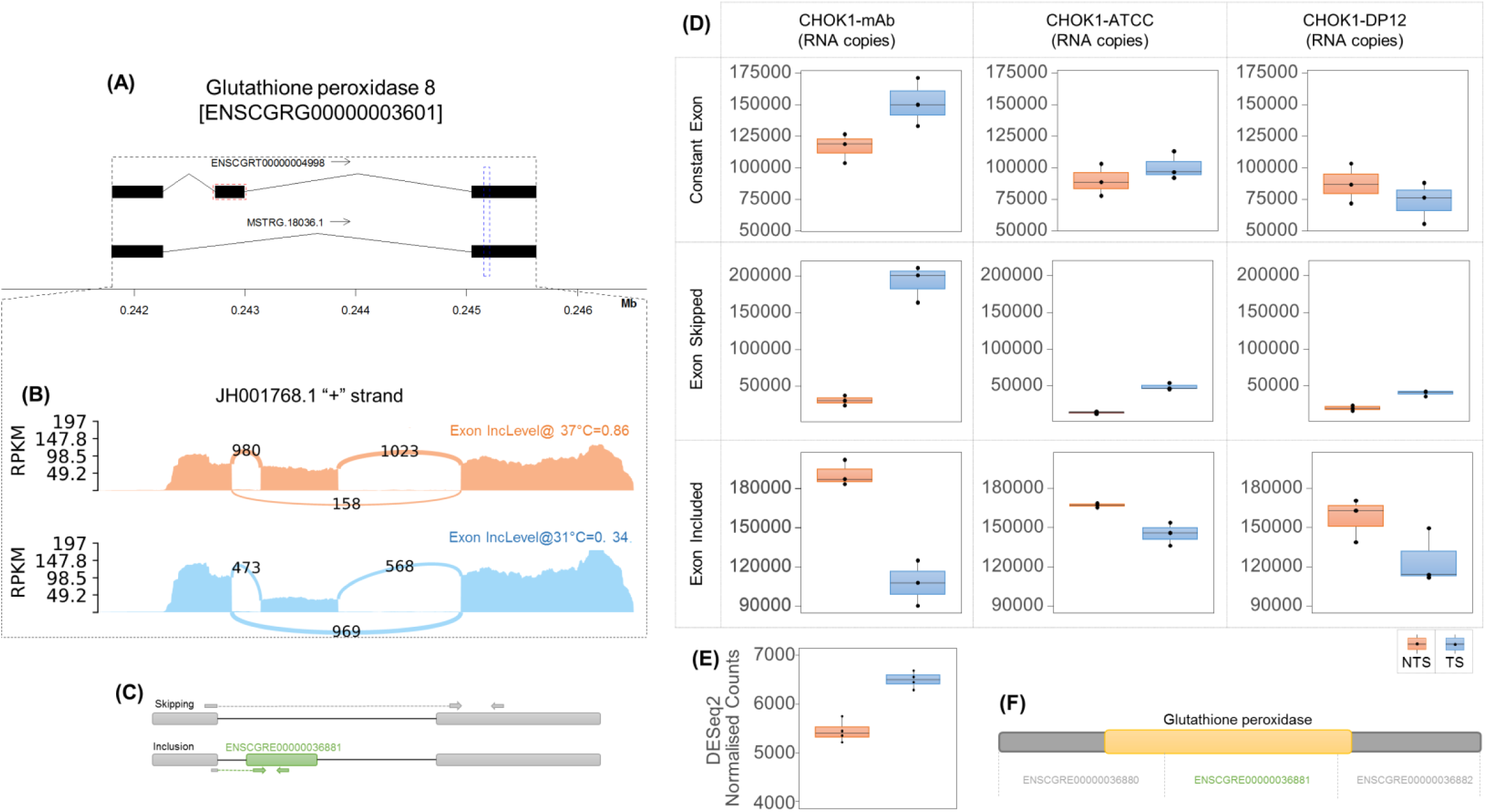
Glutathione peroxidase 8 gene is alternatively spliced following a reduction of CHO cell culture temperature. Genome-guided transcriptome assembly using the RNASeq revealed the presence of **(A)** a Gpx8 transcript not annotated in Ensembl was identified missing the second exon (ENSCGRT00000004998, highlighted in red). The annotation track was constructed using the Sushi Bioconductor package (Phanstiel, Boyle, Araya, & Snyder, 2014). The second exon of Gpx8 was found to be skipped more frequently (Δψ ≥ 50% [p = < 1 × 10^−16^]) at sub-physiological temperature following rMats analysis. A sashimi plot of the NTS and TS samples generated using rmats2sashimiplot **(B)** illustrates the read support for the *Gpx8* exon 2 skipping event from the RNASeq data (reads spanning splice junctions are shown). For qPCR confirmation of the alternative splicing event we designed **(C)** primers to quantitate the expression of isoforms including or excluding the exon as well as primers for a region of exon 3 that did not undergo alternative splicing (highlighted by blue region in **(A)**. Absolute quantitation qPCR was utilized to determine the RNA copies for NTS and TS samples from 3 cell lines. A 10% increase or decrease of RNA copies for isoforms including or excluding Gxp8 exon 2 along with a p-value of < 0.05 was considered significant. An increase in the **(D)** expression of Gpx8 transcripts without exon 2 ranging from ∼2-6 fold was observed for the 3 cell lines, while a decrease in the exon 2 included form was observed in both CHOK1 cell lines but not CHOK1-DP12 following temperature shift. For the constitutive exon no significant difference was found across the 3 cell lines in agreement with the DESeq2 RNASeq gene level expression analysis where **(E)** a ∼1.19 fold (p = 2.23 × 10^−8^) increase was found in the TS sample group. Exon 2 of *Gpx8* **(F)** is annotated in Ensembl to encode for a large portion of Glutathione peroxidase domain (Pfam ID = PF00255) for the Gpx8 protein.

Alternative splicing analysis using RNASeq revealed that the selected exon skipping events in *Dnm1l, Dnajc15* and *Mff* decreased and therefore the exon of interest was spliced in more frequently in isoforms of these genes following a reduction of cell culture temperature (Figure 4B). qPCR confirmed the inclusion of each exon of interest for the 3 genes for the two CHOK1 cell lines post-temperature shift while only the increase of the *Dnajc15* exon inclusion was confirmed for the CHOK1-DP12 cell line. A reciprocal decrease in exon skipping was observed for *Dnm1l* (Figure 6) and *Dnajc15* in the CHOK1-mAb cell line. None of the 3 reciprocal exon skipping events were found to be significantly altered in the CHOK1-ATCC cell line, while a decrease in exon skipping was identified for *Dnm1l* and *Mff* in the CHOK1-DP12 cell line.

**Figure 6:**
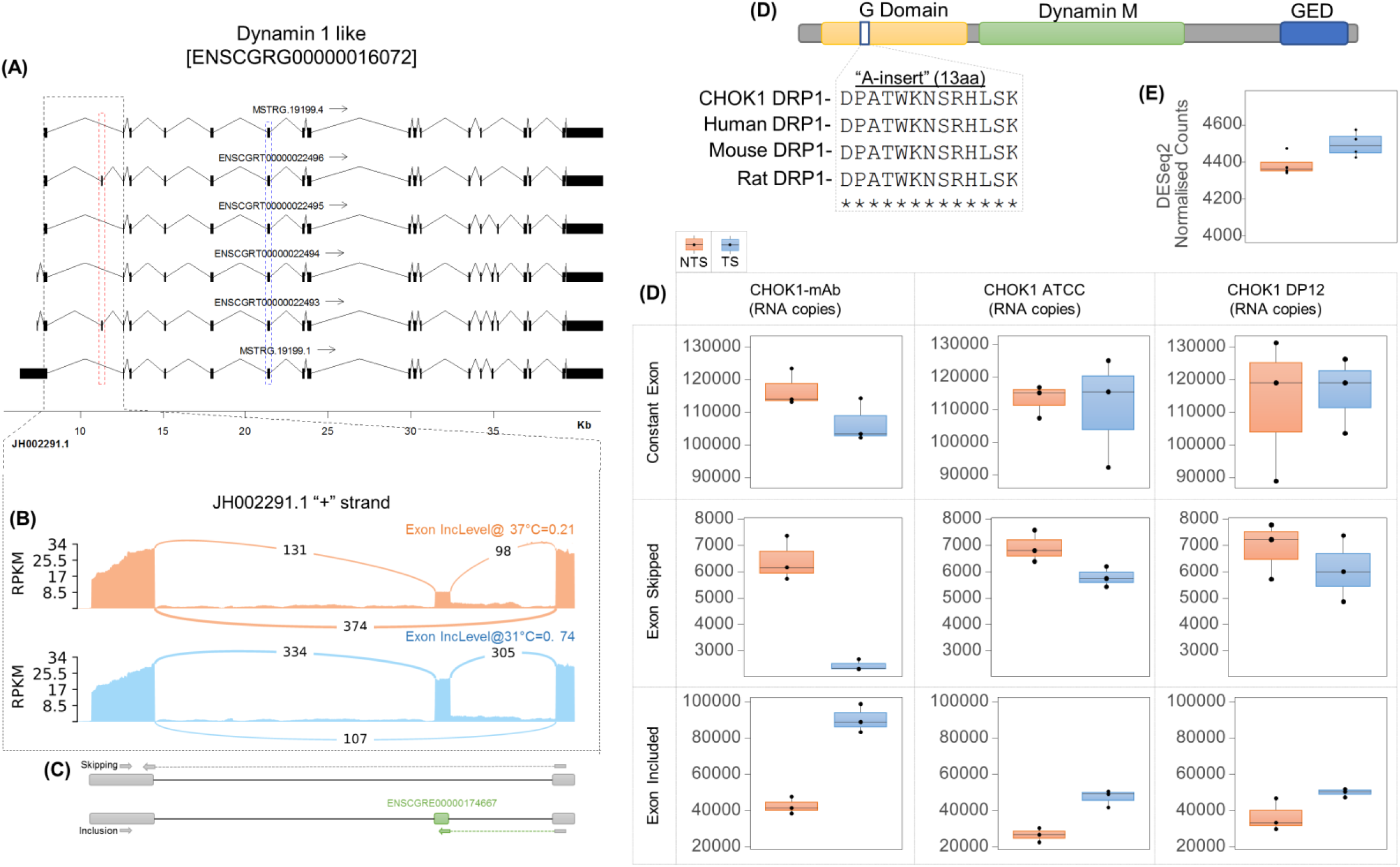
Reducing cell culture temperature results in the increased expression of *Dnm1l* isoforms containing exon 3 of the gene. Following transcriptome assembly using Stringtie an **(A)** additional 2 isoforms (MSTRG.19199.1, MSTRG.19199.4) comprised of additional 3’ UTR sequences were identified using the RNASeq data generated in this study. The annotation track was constructed using the Sushi Bioconductor package (Phanstiel et al., 2014). A single alternative splicing event was identified for the gene (highlighted in red). The sashimi plot of reads covering the splice junctions **(B)** show that a reduction of cell culture temperature results in higher inclusion of exon 3 (ENSCGRE00000174667) for Dnm1l transcripts. Primers were designed **(C)** to assess the presence of the alternate splicing event across the two conditions in the 3 cell lines. The resulting absolute qPCR **(D)** confirmed that isoforms that skipped exon 3 decreased, whereas exon3 containing isoforms increased following temperature shift in the CHOK1-mAb cell line. For the CHOK1-ATCC cell line we observed a decrease in expression of isoforms with exon 3 but no significant decrease in expression of the isoforms without exon 3. While we observed an increase for the including Dnm1l isoforms in the CHOK1-DP12 cell line was not significant. For the constitutive exon of Dnm1l (highlighted by blue region of (A)) we observed no differential expression in the 3 cell lines analysed by qPCR in agreement with the results of gene level differential expression **(E)** of the RNASeq data. Exon 3 of Dnml1 encodes for a conserved 39bp region of the Drp1 **(F)** protein called the A-insert within the G Domain of the protein annotated by Ensembl.

The primer sets for MXE splicing events in 3 genes (*Rnf34, Fars2* and *Ubr2*) were designed to detect isoforms that included one but not the other of a pair of exons (i.e. MXExon1 and MXExon2). *Rnf34* isoforms containing MXExon1 of the event were found to be increasingly expressed in all 3 cell lines post-temperature shift. A decrease in isoforms containing MXExon2 was found for CHOK1-mAb but not for CHOK1-ATCC and CHOK1-DP12. Isoforms of the *Fars2* gene containing MXExon1 were found to decrease in expression following a reduction in cell culture in the two CHOK1 cell lines, while isoforms containing Exon2 increased in expression in all three cell lines. The expression of isoforms with the inclusion of the Exon1 of the *Ubr2* MXE event decreased in all 3 cell lines, however an increase in Exon2 expression was found only for the CHOK1-ATCC cell line (Figure 4B).

## Discussion

In this manuscript, we describe the utilisation of RNASeq to characterise transcriptomics changes induced by temperature shift of a mAb producing CHOK1 cell line. For the initial phase of our analysis we sought, similar to other studies (Bedoya-López et al., 2016; Tossolini, López-Díaz, Kratje, & Prieto, 2018), to identify gene-level changes in expression following a decrease in cell culture temperature. Using a count-based differential expression method the abundance of > 1,300 genes were found to be significantly altered 24hrs after the temperature was reduced from 37°C to 31°C. In addition, *Cirbp* and *Rbm3*, well known genes involved in the cellular response to cold shock were both found to be upregulated (Masterton & Smales, 2014). Subsequent enrichment analysis highlighted the overrepresentation of genes involved in GO biological processes related to cellular proliferation and cell cycle consistent with the 24% reduction in cell number of the CHOK1-mAb cell line and previous reports in the literature (Marchant, Al-Fageeh, Underhill, Racher, & Smales, 2008).

While differential gene expression analysis is of considerable value, the principal objective of this study was to identify significant alterations in mRNA splicing and provide a deeper understanding of the impact of sub-physiological CHO cell culture temperature. The acquisition of high resolution RNASeq data (≥ 50 million 150bp paired-end reads per sample) was a crucial factor in the experimental design and enabled the accurate analysis of differential alternative splicing. The RNASeq dataset also permitted the construction of a high-quality genome guided transcriptome assembly using StringTie that greatly increased the number of CHOK1 transcripts available compared to the Ensembl annotation. The analysis of the RNASeq using the rMats pipeline when combined with the enhanced transcriptome annotation detected over 2,000 alternative splicing events where the percent spliced in value was altered by at least 10% post temperature shift. The utility of the alternative splicing analysis conducted in this study is illustrated by the identification of 1,163 protein coding genes which were unchanged in expression between the two groups yet were found to undergo 1 or more significant alternative splicing event following a reduction in cell culture temperature from 37°C to 31°C. The additional insights gained are further evidenced through the enrichment of pathways related to the metabolism of phosphate and nitrogen containing compounds identified following analysis of those genes regulated exclusively through splicing.

Of the splicing events detected exon skipping was the most frequent accounting for 50% of all the events identified, in agreement with other studies of alternative splicing (Sultan et al., 2008). Each of the 7 SE events chosen for qPCR were confirmed for CHOK1-mAb and CHOK1-ATCC cell lines, while 5 were confirmed for CHOK1-DP12. There was a lower level of agreement in SE events inferred by RNASeq (for example when an exon was found to be increasingly spliced-out by RNASeq we assessed whether the spliced-in isoform expression decreased). Changes in these inferred events were confirmed by qPCR for 5 of the SE events in CHOK1-mAb whereas only a single reciprocal event was identified in each of the other two cell lines. The differential splicing of mutually exclusive exons comprised 18.9% of all events detected. While recent evidence points to mutually exclusive exon events occurring more frequently than previously thought (Hatje et al., 2017), the possibility that the relatively early and comparatively incomplete annotation of the CHOK1 genome may have given rise to false positives or negatives cannot be ruled out. qPCR analysis of 3 MXE splicing events confirmed 5 of the 6 of the exon expression changes for the two CHOK1 cell lines and 4 of 6 events in CHOK1-DP12. The sequence conservation of human MXE splicing has been found to be greater than that of SE events highlighting their potential functional significance (Wang et al., 2008). Analysis of the *Ubr2* MXE event revealed that the two exons were found to encode a similar 33aa protein sequence (19 conserved residues) indicating these exons arose from an exon duplication event (Letunic, Copley, & Bork, 2002). The proportion of retained intron, alternative 5’ and 3’ splice site events found to be differentially spliced in this study were 8.3%, 10% and 7.6% respectively.

Considering that the RNASeq was carried out for CHOK1-mAb it is unsurprising that largest proportion of events were confirmed by qPCR in this cell line. The reasons for the weaker correlation of CHOK1-ATCC and CHOK1-DP12 to the RNASeq result, particularly with inferred events, remain unclear. It is possible that the reduction in cell culture induced a cell-line specific transcriptional response in the 24hr period was a contributory factor. Despite consistent upregulation of *Cirbp* 24hrs post-temperature shift, there was a variable reduction in cell number and the concentrations of metabolites in the cell culture media for each cell line. The different cell culture media used for CHOK1-mAb and the other two cell lines could also have played a role in the variable phenotypic responses observed for each cell line. Nevertheless, there is evidence for the occurrence of the splicing events analysed by qPCR in each of the CHO cell lines examined supporting the potential that these shared events may represent biologically significant transcriptional changes.

For instance, differential splicing was confirmed for two genes previously linked to the cellular response to environmental temperature outside of the normal physiological range. The RING finger 34 protein, another E3 ubiquitin ligase encoded by the *Rnf34* gene, has been shown to be a negative regulator of brown fat energy metabolism during the cellular response to cold through ubiquitination of PGC-1α (Wei, Pan, Mao, & Wang, 2012, p. 3). In that study, *Rnf34* gene expression was downregulated following the incubation of brown fat cells at 4°C. The results reported in this manuscript demonstrate that exon usage in *Rnf34* isoforms are altered by a reduction in temperature to mild hypothermic conditions and raise the possibility that alternative splicing may fine-tune the activity of the Rnf34 protein. Exon skipping in a second gene linked to the cellular response to temperature change, *Hikeshi*, was also confirmed by qPCR in the 3 CHO cell lines. In contrast to Rnf34, the Hikeshi protein has previously been associated with heat shock and is thought to ensure cell survival by transporting Hsp70 to the nucleus to counteract stress induced damage (Kose, Furuta, & Imamoto, 2012). While the significance of the results reported here requires further investigation, this is the first report of alternative splicing of *Hikeshi* induced by a decrease in cell culture temperature.

The difference in exon usage in the glutathione peroxidase 8, is a clear example of the impact of alternative splicing on protein sequence. The *Gpx8* gene contains 3 exons with exon 2 found to be skipped more frequently after temperature shift, encoding the majority of the glutathione peroxidase domain thought to be essential for the antioxidant activity of the protein (Nutter, Jaworski, Verma, Perez-Carrasco, & Kuyumcu-Martinez, 2017). Furthermore, a recent study has shown that the protein is enriched in mitochondria associated ER membranes and that both the transmembrane and the enzymatic domain are essential for calcium homeostasis. The remaining genes analysed by qPCR are associated with aspects of mitochondrial function including protein translation (*Slirp* & *Fars2*), activation of nuclear encoded mitochondrial proteins (*Immp1l*), regulation of metabolism (*Dnajc15*) and mitochondrial dynamics (*Dnm1l* & *Mff*).

The SRA stem-loop Interacting RNA binding protein encoded by *Slirp* has been shown to play a role in maintaining mitochondrial RNA stability as well as controlling the rate of mitochondrial protein synthesis (Lagouge et al., 2015). Reducing the cell culture temperature resulted in an increase skipping of *Slirp* exon 2 by more than 50%. Isoforms of *Slirp* lacking exon 2 alter the reading frame of the protein resulting in the introduction of a premature stop codon truncating the full-length protein by 47 residues. The *Fars2* gene, involved in transfer of phenylalanine to its cognate tRNA in mitochondria (Klipcan, Moor, Kessler, & Safro, 2009), was found to have a mutually exclusive exon splicing event was identified within its 5’ untranslated region (UTR). Approximately 13% of all human 5’ UTRs are thought to undergo splicing (Carninci et al., 2005) and it has been shown that alternative splicing can affect the translation efficiency of the resulting mRNAs. The utilisation of exons containing a upstream open reading frames can regulate the translation of the main open reading frame by interfering with ribosomal scanning (Hinnebusch, Ivanov, & Sonenberg, 2016). Increased skipping of exon 1 of the *Immp1l* gene encoding a critical protease localised to the inner mitochondrial membrane was reported to activate nuclear encoded proteins including DIABLO/Smac (Burri et al., 2005). The *Immp1l* skipped exon is unusual in that the sequence contains the final 67bp of the 5’UTR sequence and the first 105bp of the annotated open reading frame. The isoforms lacking exon 1 lose their start codon and are thus not predicted to encode for a functional protein, however translation initiation at an inframe near cognate UUG codon localised at position 252 of the shorter isoform can lead to the synthesis of a 129 aa Immp1l proteoform containing the 2 peptidase domains.

A 129bp exon of *Dnajc15* gene was found to be increasingly spliced-in by >29% post temperature shift. The Dnajc15 protein also known as MCJ is localised to the inner mitochondrial membrane and thought to be an essential negative regulator of mitochondrial metabolism (Hatle et al., 2013). The reduction of temperature shift in industry can result in a transition from an aerobic glycolysis metabolic phenotype toward the increased utilisation of oxidative phosphorylation to generate energy (Young, 2013). The Dnajc15 gene was found to be expressed before and after temperature shift and while a change in splicing of Dnajc15 upon a reduction of temperature may, at first, seem counter intuitive, it has been estimated that OXPHOS can also contribute significantly to ATP production in parallel with aerobic glycolysis during exponential growth (Young, 2013). While further studies are required, the alternative splicing in Dnajc15 could play a role in negative regulation of mitochondrial metabolism upon growth arrest.

The traditional view of single, stationary mitochondria has been replaced by understanding that these organelles are in fact dynamic, forming interconnected networks and undergoing quality control through coordinated rounds of fission and fusion. Temperature shift induced alternative splicing of the *Dnml1* and *Mff* genes encoding dynamin related protein 1 (Drp1) and mitochondrial fission factor (Mff), two critical regulators of mitochondrial dynamics (Tilokani, Nagashima, Paupe, & Prudent, 2018). Drp1 is a member of the dynamin GTPase family and is recruited by outer mitochondrial membrane bound Mff. Drp1 monomers oligomerize around the mitochondria before constricting the mitochondrial membrane in a GTP dependent manner. The *Dnm1l* gene is expressed as multiple isoforms in mammalian cells through alternative splicing of exons 3, 16 and 17. These isoforms encode the variable regions of Drp1 known as the A-insert (exon 3) and B-insert (exons 16 & 17) (Macdonald et al., 2016). Upon the reduction of cell culture temperature, the inclusion of exon 3 increased in *Dnm1l* isoforms but no significant splicing difference was found for the other two exons. The A-insert encodes a 13aa protein sequence that has been shown to inhibit Drp1 GTPase activity in conjunction with the B-insert. The interaction of Drp1 with Mff is thought to alleviate this inhibition of Drp1 in an allosteric manner. Similar to Drp1 qPCR confirmed that an *Mff* exon was included more frequently post temperature shift, however considering that 3 SE, 2 MXE and a RI event were also identified by RNASeq the impact of a single event is, at this point, unclear. While the alternative splicing patterns of the *Mff* gene are highly complex and require and further analysis is required to determine the biological significance of these findings, the results of this study suggest that the regulation of mitochondrial dynamics via alternative splicing may play an important role in the CHO cell response to a temperature shift.

## Conclusions

The findings of this study reveal that the transition of CHO cells to a sub-physiological bioreactor temperature induces a transcriptional response more complex than previously appreciated. The cellular response to temperature shift not only results in the widespread alteration of gene expression but also has an equally dramatic effect on mRNA splicing. Indeed, many of the genes found to be alternatively spliced would have been ignored had differential gene expression analysis been used in isolation. For instance, isoforms arising from several genes involved in mitochondrial function were confirmed to undergo alternative splicing yet no significant difference in gene expression was found. The utilisation of RNASeq based analysis of alternative splicing provides an additional dimension to our understanding of transcriptional regulation and will be a valuable tool in future studies of CHO cell biology.

## Acknowledgments

The authors gratefully acknowledge funding from Science Foundation Ireland (grant references 13/SIRG/2084 & 15/CDA/3529) and H2020 Marie Sklodowska-Curie (Grant agreement No: 642663 & No: 813453).

## Supplementary Figures

**Figure S1:**
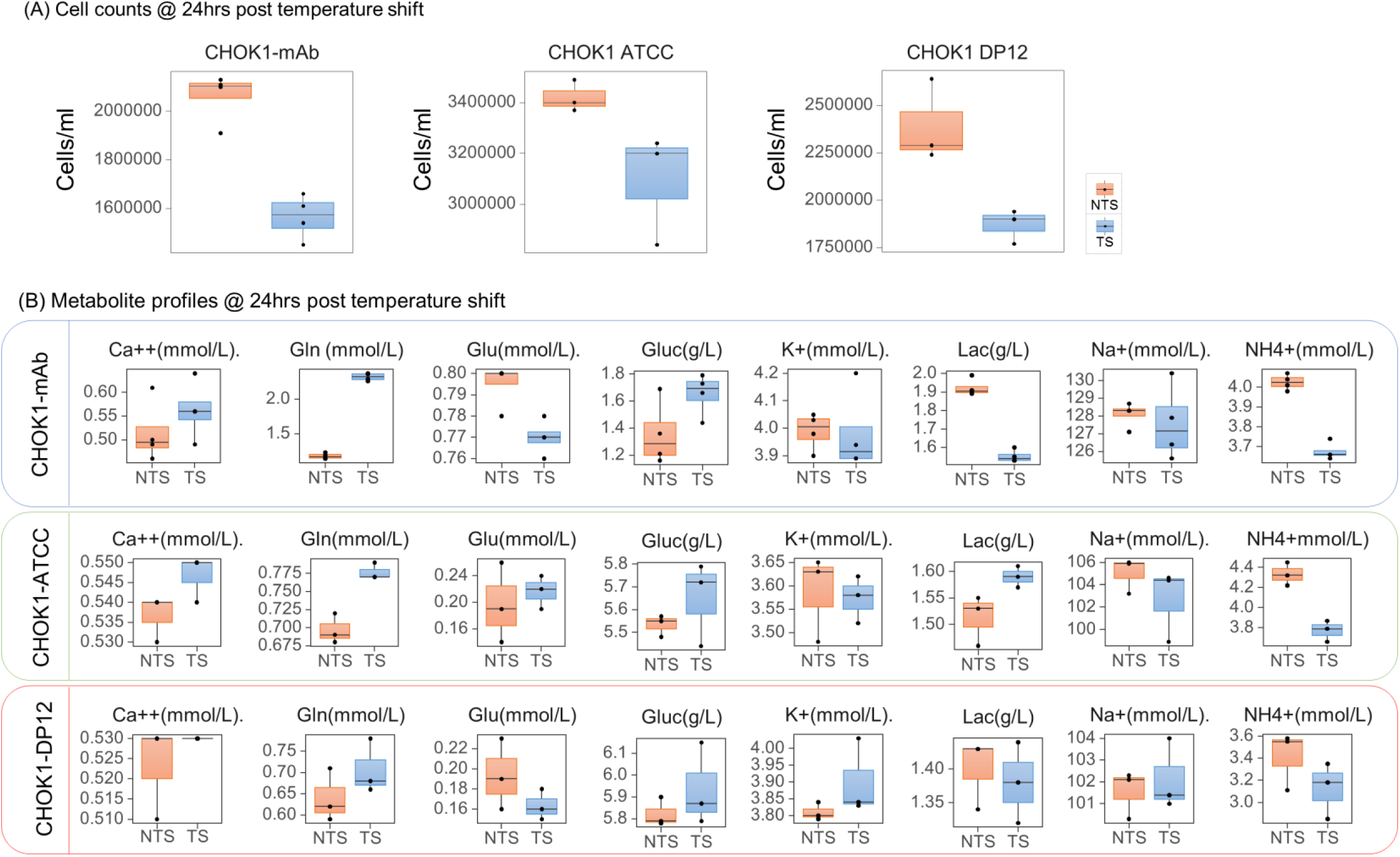
**(A)** Cell counts as well as for the CHOK1-ATCC and CHOK1-DP12 in the TS and NTS sample groups as well as **(B)** Nova Flex analysis of calcium, lactate, ammonium, sodium, potassium, glutamine, glutamate and glucose for the CHOK1-mAb, CHOK1-ATCC and CHOK1-DP12 cell lines.

**Figure S2:**
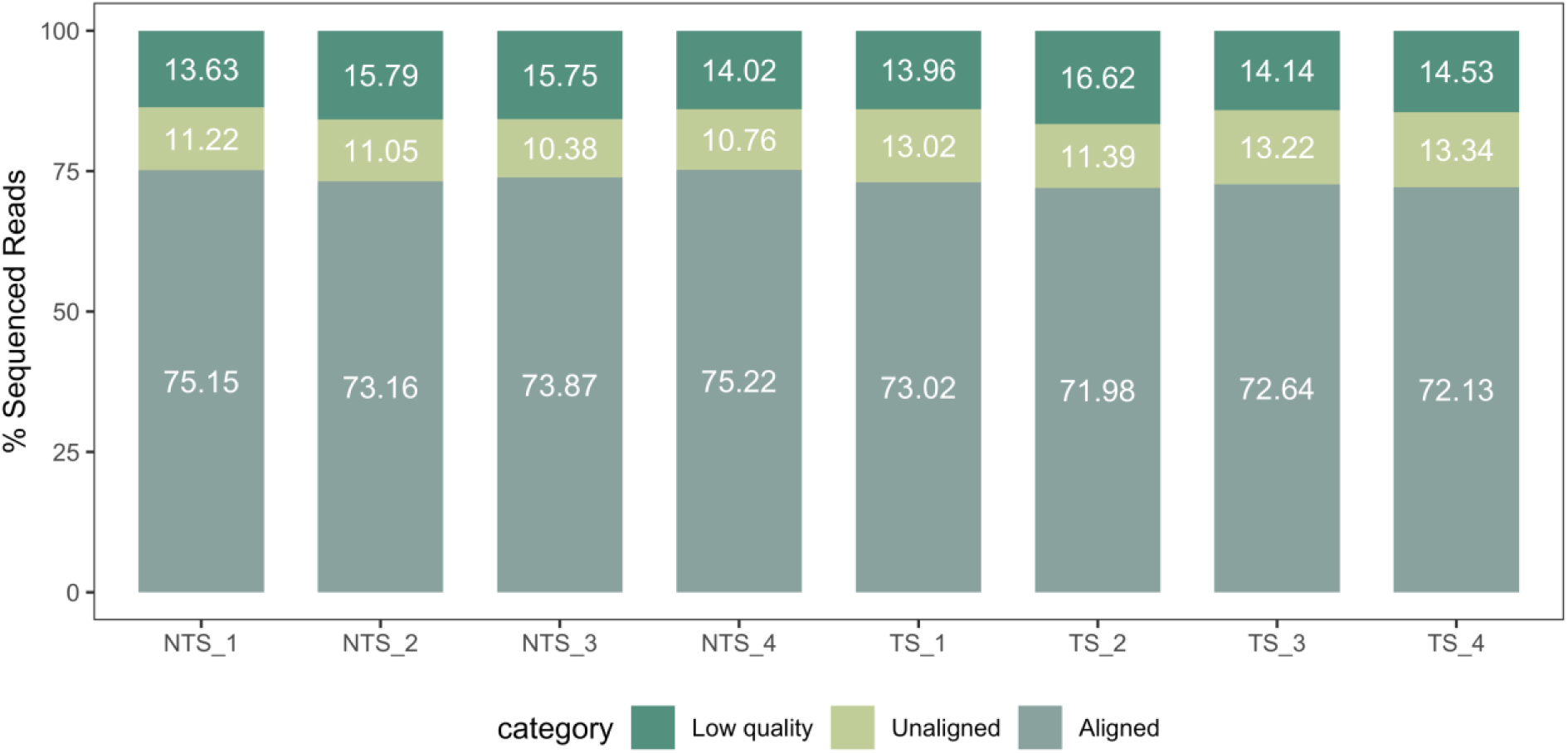
Proportion of reads from each sample that were removed following preprocessing and those that aligned to the genome.

**Figure S3:**
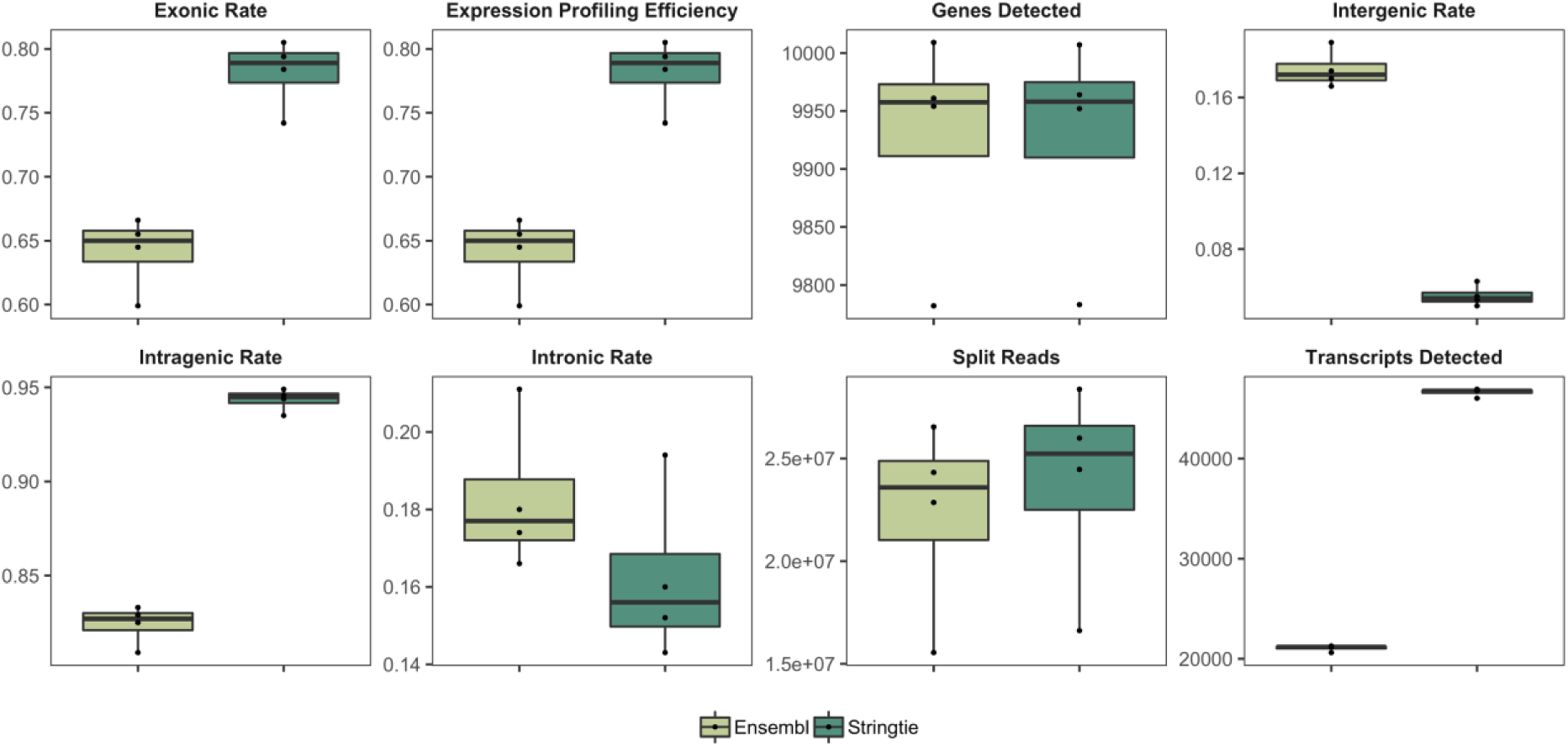
RNASeQC analysis of the effectiveness of the RNASeq library preparation, alignment and genome guided assembly using StringTie.

**Figure S4:** Standard curves for absolute qPCR of *Cirbp* and selected alternative splicing events. **Download link:** https://app.box.com/s/bufhwz17abrqc92ej3ytqvt5cgmebo1v

## Supplementary Tables

**Table S1: CHOK1-mAb cell culture characteristics for the 3 CHO cell lines. (A)** CHOK1-mAb cell counts for the NTS and TS samples at 24, 48 and 72hrs post-seeding. **(B)** CHOK1-mAb Nova Flex extracellular metabolites at 72hrs post-seeding. **(C)** CHOK1-ATCC cell counts at 72hrs post-seeding. **(D)** CHOK1-ATCC Nova Flex extracellular metabolite at 72hrs post-seeding. **(E)** CHOK1-DP12 cell counts at 72hrs post-seeding. **(F)** CHOK1-DP12 Nova Flex extracellular metabolite at 72hrs post-seeding. **Download link:** https://app.box.com/s/5qn2rurbhnl1n3r8fze6yujrrei0gegs

**Table S2: RNASeq read and alignment statistics (A)** The total number of raw sequencing reads acquired per sample are shown along with the number of reads remaining after processing and the number of reads aligned to the genome. **Download link:** https://app.box.com/s/w6aa3kqkkzcb3tt2qxd7pdwxvffxe80f

**Table S3: Differentially expressed gene list**. 1317 genes (1278 annotated in either ENSEMBL or NCBI) were found to be differentially expressed upon comparison of the NTS and TS sample groups. The ENSEBML ID, NCBI ID, gene symbol, Gene Description, baseMean of DESeq2 normalised counts, log2 p-value and BH adjusted p-value are shown for each gene. **Download link:** https://app.box.com/s/dihyod0yrbkbfzysonol8nfh1wvjvds4

**Table S4: Differential splicing genelist**. Statistically significant alternative splicing events were identified by RNASeq for CHOK1 cell line separated into **(A)** Skipped exons, **(B)** Mutually exclusive exons, **(C)** Retained introns as well as alternative **(D)** 5’ and **(E)** 3’ splice sites. **Download link:** https://app.box.com/s/mrljl4c0kq6nw23b9r5ke6dys5igdop1

**Table S5: GOrilla enrichment analysis** for the **(A)** AS genes GO biological processes **(B)** AS GO genes molecular functions **(C)** AS GO genes cellular compartment **(D)** DE genes GO biological processes **(E)** DE genes GO molecular functions **(F)** DE genes GO cellular compartment. **Download link:** https://app.box.com/s/4u0ms0xsuoxf2uqirnk74tjctlaf9t34

**Table S6:** Primer design for Cirbp, selected alternative splicing events and constitutive exons. **Download link:** https://app.box.com/s/rs559g9ua5uk1s8cz4uu1mh9qct9fpk9

**Table S7:** Gblock sequence design for absolute qPCR. **Download link:** https://app.box.com/s/fhqkhml4sp6xe3juii896jts8gyvgb99

**Table S8** qPCR analysis including the raw data and fold changes for the alternative splicing events, constitutive exons and *Cirbp*. **Download link:** https://app.box.com/s/h5i8yc579pxj2femcxh6uspks2rpxem9

## References

Bedoya-López, A., Estrada, K., Sanchez-Flores, A., Ramírez, O. T., Altamirano, C., Segovia, L., … Valdez-Cruz, N. A. (2016). Effect of Temperature Downshift on the Transcriptomic Responses of Chinese Hamster Ovary Cells Using Recombinant Human Tissue Plasminogen Activator Production Culture. PloS One, 11(3), e0151529. https://doi.org/10.1371/journal.pone.0151529

Boise, L. H., González-García, M., Postema, C. E., Ding, L., Lindsten, T., Turka, L. A., … Thompson, C. B. (1993). Bcl-x, a bcl-2-related gene that functions as a dominant regulator of apoptotic cell death. Cell, 74(4), 597–608.

Bolger, A. M., Lohse, M., & Usadel, B. (2014). Trimmomatic: A flexible trimmer for Illumina sequence data. Bioinformatics, 30(15), 2114–2120. https://doi.org/10.1093/bioinformatics/btu170

Brinkrolf, K., Rupp, O., Laux, H., Kollin, F., Ernst, W., Linke, B., … Borth, N. (2013). Chinese hamster genome sequenced from sorted chromosomes. Nature Biotechnology, 31(8), 694–695. https://doi.org/10.1038/nbt.2645

Burri, L., Strahm, Y., Hawkins, C. J., Gentle, I. E., Puryer, M. A., Verhagen, A., … Lithgow, T. (2005). Mature DIABLO/Smac is produced by the IMP protease complex on the mitochondrial inner membrane. Molecular Biology of the Cell, 16(6), 2926–2933. https://doi.org/10.1091/mbc.e04-12-1086

Carninci, P., Kasukawa, T., Katayama, S., Gough, J., Frith, M. C., Maeda, N., … Hayashizaki, Y. (2005). The Transcriptional Landscape of the Mammalian Genome. Science, 309(5740), 1559–1563. https://doi.org/10.1126/science.1112014

Clarke, C., Gallagher, C., Kelly, R. M., Henry, M., Meleady, P., Frye, C. C., … Clynes, M. (2019). Transcriptomic analysis of IgG4 Fc-fusion protein degradation in a panel of clonally-derived CHO cell lines using RNASeq. Biotechnology and Bioengineering. https://doi.org/10.1002/bit.26958

DeLuca, D. S., Levin, J. Z., Sivachenko, A., Fennell, T., Nazaire, M.-D., Williams, C., … Getz, G. (2012). RNA-SeQC: RNA-seq metrics for quality control and process optimization. Bioinformatics (Oxford, England), 28(11), 1530–1532. https://doi.org/10.1093/bioinformatics/bts196

Dobin, A., Davis, C. A., Schlesinger, F., Drenkow, J., Zaleski, C., Jha, S., … Gingeras, T. R. (2013). STAR: Ultrafast universal RNA-seq aligner. Bioinformatics (Oxford, England), 29(1), 15–21. https://doi.org/10.1093/bioinformatics/bts635

Eden, E., Navon, R., Steinfeld, I., Lipson, D., & Yakhini, Z. (2009). GOrilla: A tool for discovery and visualization of enriched GO terms in ranked gene lists. BMC Bioinformatics, 10(1), 48. https://doi.org/10.1186/1471-2105-10-48

Fischer, S., Handrick, R., & Otte, K. (2015). The art of CHO cell engineering: A comprehensive retrospect and future perspectives. Biotechnology Advances, 33(8), 1878–1896. https://doi.org/10.1016/j.biotechadv.2015.10.015

Ge, Y., & Porse, B. T. (2014). The functional consequences of intron retention: Alternative splicing coupled to NMD as a regulator of gene expression. BioEssays: News and Reviews in Molecular, Cellular and Developmental Biology, 36(3), 236–243. https://doi.org/10.1002/bies.201300156

Gerstl, M. P., Hackl, M., Graf, A. B., Borth, N., & Grillari, J. (2013). Prediction of transcribed PIWI-interacting RNAs from CHO RNAseq data. Journal of Biotechnology, 166(1–2), 51–57. https://doi.org/10.1016/j.jbiotec.2013.04.010

Hackl, M., Jadhav, V., Jakobi, T., Rupp, O., Brinkrolf, K., Goesmann, A., … Grillari, J. (2012). Computational identification of microRNA gene loci and precursor microRNA sequences in CHO cell lines. Journal of Biotechnology, 158(3), 151–155. https://doi.org/10.1016/j.jbiotec.2012.01.019

Hartley, S. W., & Mullikin, J. C. (2016). Detection and visualization of differential splicing in RNA-Seq data with JunctionSeq. Nucleic Acids Research, 44(15), e127. https://doi.org/10.1093/nar/gkw501

Hatje, K., Rahman, R.-U., Vidal, R. O., Simm, D., Hammesfahr, B., Bansal, V., … Kollmar, M. (2017). The landscape of human mutually exclusive splicing. Molecular Systems Biology, 13(12), 959. https://doi.org/10.15252/msb.20177728

Hatle, K. M., Gummadidala, P., Navasa, N., Bernardo, E., Dodge, J., Silverstrim, B., … Rincon, M. (2013). MCJ/DnaJC 15, an Endogenous Mitochondrial Repressor of the Respiratory Chain That Controls Metabolic Alterations. Molecular and Cellular Biology, 33(11), 2302–2314. https://doi.org/10.1128/MCB.00189-13

Hinnebusch, A. G., Ivanov, I. P., & Sonenberg, N. (2016). Translational control by 5’-untranslated regions of eukaryotic mRNAs. Science (New York, N.Y.), 352(6292), 1413–1416. https://doi.org/10.1126/science.aad9868

Kaas, C. S., Kristensen, C., Betenbaugh, M. J., & Andersen, M. R. (2015). Sequencing the CHO DXB11 genome reveals regional variations in genomic stability and haploidy. BMC Genomics, 16, 160. https://doi.org/10.1186/s12864-015-1391-x

Kelemen, O., Convertini, P., Zhang, Z., Wen, Y., Shen, M., Falaleeva, M., & Stamm, S. (2013). Function of alternative splicing. Gene, 514(1), 1–30. https://doi.org/10.1016/j.gene.2012.07.083

Kelly, P. S., Clarke, C., Costello, A., Monger, C., Meiller, J., Dhiman, H., … Barron, N. (2017). Ultra-deep next generation mitochondrial genome sequencing reveals widespread heteroplasmy in Chinese hamster ovary cells. Metabolic Engineering, 41, 11–22. https://doi.org/10.1016/j.ymben.2017.02.001

Klipcan, L., Moor, N., Kessler, N., & Safro, M. G. (2009). Eukaryotic cytosolic and mitochondrial phenylalanyl-tRNA synthetases catalyze the charging of tRNA with the meta-tyrosine. Proceedings of the National Academy of Sciences of the United States of America, 106(27), 11045–11048. https://doi.org/10.1073/pnas.0905212106

Kose, S., Furuta, M., & Imamoto, N. (2012). Hikeshi, a Nuclear Import Carrier for Hsp70s, Protects Cells from Heat Shock-Induced Nuclear Damage. Cell, 149(3), 578–589. https://doi.org/10.1016/j.cell.2012.02.058

Kuo, C.-C., Chiang, A. W., Shamie, I., Samoudi, M., Gutierrez, J. M., & Lewis, N. E. (2018). The emerging role of systems biology for engineering protein production in CHO cells. Current Opinion in Biotechnology, 51, 64–69. https://doi.org/10.1016/j.copbio.2017.11.015

Lagouge, M., Mourier, A., Lee, H. J., Spåhr, H., Wai, T., Kukat, C., … Larsson, N.-G. (2015). SLIRP Regulates the Rate of Mitochondrial Protein Synthesis and Protects LRPPRC from Degradation. PLOS Genetics, 11(8), e1005423. https://doi.org/10.1371/journal.pgen.1005423

Letunic, I., Copley, R. R., & Bork, P. (2002). Common exon duplication in animals and its role in alternative splicing. Human Molecular Genetics, 11(13), 1561–1567. https://doi.org/10.1093/hmg/11.13.1561

Lévêque, C., Marsaud, V., Renoir, J.-M., & Sola, B. (2007). Alternative cyclin D1 forms a and b have different biological functions in the cell cycle of B lymphocytes. Experimental Cell Research, 313(12), 2719–2729. https://doi.org/10.1016/j.yexcr.2007.04.018

Lewis, N. E., Liu, X., Li, Y., Nagarajan, H., Yerganian, G., O’Brien, E., … Palsson, B. O. (2013). Genomic landscapes of Chinese hamster ovary cell lines as revealed by the Cricetulus griseus draft genome. Nature Biotechnology, 31(8), 759–765. https://doi.org/10.1038/nbt.2624

Macdonald, P. J., Francy, C. A., Stepanyants, N., Lehman, L., Baglio, A., Mears, J. A., … Ramachandran, R. (2016). Distinct Splice Variants of Dynamin-related Protein 1 Differentially Utilize Mitochondrial Fission Factor as an Effector of Cooperative GTPase Activity. Journal of Biological Chemistry, 291(1), 493–507. https://doi.org/10.1074/jbc.M115.680181

Marchant, R. J., Al-Fageeh, M. B., Underhill, M. F., Racher, A. J., & Smales, C. M. (2008). Metabolic rates, growth phase, and mRNA levels influence cell-specific antibody production levels from in vitro-cultured mammalian cells at sub-physiological temperatures. Molecular Biotechnology, 39(1), 69–77. https://doi.org/10.1007/s12033-008-9032-0

Martin, M. (2011). Cutadapt removes adapter sequences from high-throughput sequencing reads. EMBnet.Journal, 17(1), 10–12. https://doi.org/10.14806/ej.17.1.200

Masterton, R. J., & Smales, C. M. (2014). The impact of process temperature on mammalian cell lines and the implications for the production of recombinant proteins in CHO cells. Pharmaceutical Bioprocessing, 2, 49–61.

Monger, C., Kelly, P. S., Gallagher, C., Clynes, M., Barron, N., & Clarke, C. (2015). Towards next generation CHO cell biology: Bioinformatics methods for RNA-Seq-based expression profiling. Biotechnology Journal, 10(7), 950–966. https://doi.org/10.1002/biot.201500107

Nutter, C. A., Jaworski, E., Verma, S. K., Perez-Carrasco, Y., & Kuyumcu-Martinez, M. N. (2017). Developmentally regulated alternative splicing is perturbed in type 1 diabetic skeletal muscle. Muscle & Nerve, 56(4), 744–749. https://doi.org/10.1002/mus.25599

Pan, Q., Shai, O., Lee, L. J., Frey, B. J., & Blencowe, B. J. (2008). Deep surveying of alternative splicing complexity in the human transcriptome by high-throughput sequencing. Nature Genetics, 40(12), 1413–1415. https://doi.org/10.1038/ng.259

Pertea, M., Pertea, G. M., Antonescu, C. M., Chang, T.-C., Mendell, J. T., & Salzberg, S. L. (2015). StringTie enables improved reconstruction of a transcriptome from RNA-seq reads. Nature Biotechnology, 33(3), 290–295. https://doi.org/10.1038/nbt.3122

Phanstiel, D. H., Boyle, A. P., Araya, C. L., & Snyder, M. P. (2014). Sushi.R: Flexible, quantitative and integrative genomic visualizations for publication-quality multi-panel figures. Bioinformatics, 30(19), 2808–2810. https://doi.org/10.1093/bioinformatics/btu379

Raab, N., Mathias, S., Alt, K., Handrick, R., Fischer, S., Schmieder, V., … Otte, K. (2019). CRISPR/Cas9-mediated knockout of microRNA-744 improves antibody titer of CHO production cell lines. Biotechnology Journal, e1800477. https://doi.org/10.1002/biot.201800477

Rupp, O., MacDonald, M. L., Li, S., Dhiman, H., Polson, S., Griep, S., … Lee, K. H. (2018). A reference genome of the Chinese hamster based on a hybrid assembly strategy. Biotechnology and Bioengineering, 115(8), 2087–2100. https://doi.org/10.1002/bit.26722

Sha, S., Bhatia, H., & Yoon, S. (2018). An RNA-seq based transcriptomic investigation into the productivity and growth variants with Chinese hamster ovary cells. Journal of Biotechnology, 271, 37–46. https://doi.org/10.1016/j.jbiotec.2018.02.008

Shen, S., Park, J. W., Lu, Z., Lin, L., Henry, M. D., Wu, Y. N., … Xing, Y. (2014). rMATS: Robust and flexible detection of differential alternative splicing from replicate RNA-Seq data. Proceedings of the National Academy of Sciences of the United States of America, 111(51), E5593–5601. https://doi.org/10.1073/pnas.1419161111

Sultan, M., Schulz, M. H., Richard, H., Magen, A., Klingenhoff, A., Scherf, M., … Yaspo, M.-L. (2008). A Global View of Gene Activity and Alternative Splicing by Deep Sequencing of the Human Transcriptome. Science, 321(5891), 956–960. https://doi.org/10.1126/science.1160342

Tilokani, L., Nagashima, S., Paupe, V., & Prudent, J. (2018). Mitochondrial dynamics: Overview of molecular mechanisms. Essays In Biochemistry, 62(3), 341–360. https://doi.org/10.1042/EBC20170104

Tossolini, I., López-Díaz, F. J., Kratje, R., & Prieto, C. C. (2018). Characterization of cellular states of CHO-K1 suspension cell culture through cell cycle and RNA-sequencing profiling. Journal of Biotechnology, 286, 56–67. https://doi.org/10.1016/j.jbiotec.2018.09.007

Walsh, G. (2018). Biopharmaceutical benchmarks 2018. Nature Biotechnology, 36, 1136–1145. https://doi.org/10.1038/nbt.4305

Wang, E. T., Sandberg, R., Luo, S., Khrebtukova, I., Zhang, L., Mayr, C., … Burge, C. B. (2008). Alternative isoform regulation in human tissue transcriptomes. Nature, 456(7221), 470–476. https://doi.org/10.1038/nature07509

Wei, P., Pan, D., Mao, C., & Wang, Y.-X. (2012). RNF34 is a cold-regulated E3 ubiquitin ligase for PGC-1α and modulates brown fat cell metabolism. Molecular and Cellular Biology, 32(2), 266–275. https://doi.org/10.1128/MCB.05674-11

Will, C. L., & Lührmann, R. (2011). Spliceosome Structure and Function. Cold Spring Harbor Perspectives in Biology, 3(7). https://doi.org/10.1101/cshperspect.a003707

Xu, X., Nagarajan, H., Lewis, N. E., Pan, S., Cai, Z., Liu, X., … Wang, J. (2011). The genomic sequence of the Chinese hamster ovary (CHO)-K1 cell line. Nature Biotechnology, 29(8), 735–741. https://doi.org/10.1038/nbt.1932

Yoshida, H., Matsui, T., Yamamoto, A., Okada, T., & Mori, K. (2001). XBP1 mRNA Is Induced by ATF6 and Spliced by IRE1 in Response to ER Stress to Produce a Highly Active Transcription Factor. Cell, 107(7), 881–891. https://doi.org/10.1016/S0092-8674(01)00611-0

Young, J. D. (2013). Metabolic flux rewiring in mammalian cell cultures. Current Opinion in Biotechnology, 24(6). https://doi.org/10.1016/j.copbio.2013.04.016

